# Genetic diversity of domestic cat hepadnavirus in Taiwan

**DOI:** 10.1101/2023.09.29.560097

**Authors:** Benji Brayan Ilagan Silva, Jin-Yang Chen, Brian Harvey Avanceña Villanueva, Zi-Ying Lu, Hsing-Hua Zhen, Andrew D. Montecillo, Maya Shofa, Hoang Minh, Jen-Pin Chuang, Huai-Ying Huang, Akatsuki Saito, Kuo-Pin Chuang

## Abstract

Domestic cat hepadnavirus (DCH) is an infectious disease associated with chronic hepatitis in cats, suggesting a similarity with hepatitis B virus infection in humans. Since its first identification in Australia in 2018, DCH has been reported in several countries with varying prevalence rates, but its prevalence in Taiwan has not yet been investigated. Here, we aimed to identify the presence and prevalence of DCH infections in Taiwan. Among 71 samples tested, eight (11.27%) were positive for DCH. Of these positive cases, three cats had elevated levels of alanine transaminase (ALT) and aspartate transaminase (AST), suggesting an association between DCH infection and chronic hepatitis. Four DCH-positive samples were also tested for feline immunodeficiency virus (FIV) and feline leukemia virus (FeLV) co-infection, one (25%) was positive for FIV while none for FeLV (0%). In addition, we performed whole genome sequencing of six samples to determine the viral genome sequences. Phylogenetic analyses identified a distinct lineage compared with previously reported sequences. Considering the recent findings suggesting the potential risk of DCH for interspecies or zoonotic transmission, this study suggests the importance of continuous surveillance of DCH and further research to elucidate the pathophysiology and transmission route of DCH.

## 1. Introduction

Domestic cat hepadnavirus (DCH), a novel member of the family *Hepadnaviridae* belonging to the genus *Orthohepadnavirus*, was recently identified through a virus discovery transcriptomic study on a large cell lymphoma collected from an immunocompromised FIV-positive domestic cat with hepatic disease [1]. Like other members of the genus *Orthohepadnavirus*, DCH contains partially double-stranded, relaxed circular DNA molecule of about 3.2 kb in length enclosed in an envelope. The genome has four overlapping gene regions encoding the polymerase, surface, core, and X proteins [1].

Previous studies have shown a significant frequency of DCH detection in cats suffering from chronic hepatitis and hepatocellular carcinoma, like HBV infection in humans. Therefore, it is crucial to conduct further investigations to establish a direct association between DCH infection and hepatitis in cats. One potential approach for validation involves employing *in situ* hybridization (ISH) to confirm the presence of DCH in regions of liver inflammation [2]. Moreover, the presence of DCH was also identified in various organs other than the liver, including the heart, lungs, intestines, kidneys, and spleen pointing to its broad cellular tropism [1,3–6]. On the other hand, repeated testing of cat**s** naturally infected with the DCH showed negative PCR assay results for oral, conjunctival, preputial, and rectal swabs. However, the detection of the DCH in the blood and intestinal samples in the same study and in other previous studies suggests that the virus may be transmitted via blood or fecal material [3,6,7].

Following its initial discovery in Australia, investigation of DCH prevalence of this virus in cats from different countries have been carried out, with positive cases reported in Italy, Thailand, Malaysia, the United Kingdom, Japan, the United States, Hong Kong, and Türkiye [1–17]. However, there is a lack of information regarding this virus in Taiwan, highlighting the importance of DCH investigation.

## 2. Materials and Methods

### 2.1 Ethics Statement

All experiments in this study followed the protocols and guidelines approved by the Institutional Animal Care and Use Committee of the National Pingtung University of Science and Technology (NPUST-IACUC), with approval number NPUST-112-079.

### 2.2 Sample Collection

Cat blood samples were obtained through multiple clinics in southern Taiwan following the collection and sample handling guidelines approved by the relevant committee as mentioned above. Residual or leftover samples from DCH-positive cats were used to test for liver health markers including alanine transaminase (ALT), aspartate transaminase (AST), and alkaline phosphatase (ALKP). DCH-positive whole blood residue samples were also tested for the FIV and FeLV co-infection using a SNAP^®^ FIV/FeLV Combo Test (IDEXX, Taiwan, 99-08354). DCH-positive patients were invited for a follow-up health check 2.5 months after initial detection to monitor changes in viral loads and liver health markers.

### 2.3 DCH detection by direct duplex real-time quantitative PCR

Detection of the domestic cat hepadnavirus was performed as described previously [18], with minute modifications following the manufacturer’s instructions on the use of Premix Ex Taq™ (Probe qPCR) mix (TaKaRa, Japan, RR390L). Briefly, 1 μL of whole blood sample was added into the PCR reaction mix containing 0.2 μL of each of the primer (forward and reverse), 0.1 μL of each of the probes at 10 μM concentrations, 10 μL of 2× Probe qPCR mix, 0.4 μL of ROX reference dye, and 7.6 μL double-distilled sterilized PCR-grade water for a total volume of 20 μL. The reaction was performed using a StepOne™ realtime thermal cycler system (Applied Biosystems, Singapore). All primers, probes, and reaction conditions were as described elsewhere [18].

### 2.4 PCR Amplification of DCH whole genome

Total genomic DNA from the serum samples that tested positive for DCH was extracted using Viogene (Blood and Tissue Genomic Mini) (Viogene, Taiwan, GG1001) following the manufacturer’s Blood Protocol. Three sets of primers derived from [10] were used to amplify overlapping fragments covering the whole genome of DCH (Table 1). The PCR reaction, in a final volume of 50 μL, contained 25 μL P Easy-Pfu 2X PCR SuperMix (AllBio, Taiwan, ABTGMBP03-100), 1 μL of each primer (forward and reverse) at 10 μM concentrations, 10 μL extracted DNA, and 13 μL double-distilled sterilized PCR-grade water. Amplification conditions were as follows: initial denaturation at 94°C for 5mins, followed by 40 cycles of 94°C for 30 sec, 53°C for 30 sec, and 72°C for 3min and 30 sec, followed by a final extension at 72°C for 10 min. Amplicons were purified to remove the unused enzymes and dNTPs using PCR Clean-Up & Gel Extraction Kit (Bio-Helix, Taiwan, PDC01-0100) following the PCR Cleanup protocol as directed by the manufacturer.

**Table 1.**
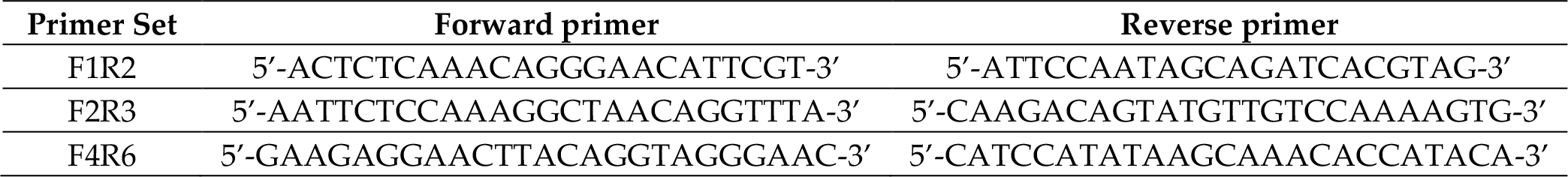
Primers used for the amplification of the DCH whole genome.

### 2.5 Sequencing and Assembly

Purified PCR amplicons were barcoded using the SQK-RBK114.24 kit (Oxford Nanopore Technologies, UK, SQK-RBK114.24) and loaded on a MinION SpotON R10.4.1 FLO-MIN114 flow cell (Oxford Nanopore Technologies, UK, R10.4.1 FLO-MIN114). The sequencing was run for 7 hours in MinKNOW Software (v.23.04.6) on a MinION MK1b device. Live basecalling was performed using the super accurate, 400bps model with default parameters in Guppy v.6.5.7 (Oxford Nanopore Technologies, UK).

For each of the barcodes, basecalled reads in the fastq_pass folder were combined in one fastq file and were mapped using bwa v.0.7.17-r1188 against DCH strain TR-404 genome (GenBank: OQ130245.1). Following this, sequence quality control was performed using Porechop v0.2.4 (https://github.com/rrwick/Porechop) with default parameters (-discard_middle). Cap3 (VersionDate: 02/10/15) was used to perform the assembly, and manual curation of the assembled contigs was performed using aligned selected contigs. Alignment was performed in UGene (v.48.0) using ClustalW, allowing for a default gap opening penalty (15.00). The assembly was polished in Medaka v1.8.0 (https://github.com/nanoporetech/medaka).

The DCH genomes from Taiwan strains identified in this study were deposited in GenBank (accession no. OR515499-OR515504). The raw reads have been deposited in the NCBI Sequence Read Archive (SRA accession no. SRX21620583-SRX21620588). The BioProject and BioSample accession numbers are PRJNA1011820, and SAMN37224405 to SAMN37224410, respectively.

### 2.6 Phylogenetic and Recombination Analyses

Alignments of the recovered genome and protein sequences against available DCH and other hepadnavirus sequences were performed using Clustal (v. 2.1). Maximum likelihood trees were constructed in IQTREE v.2.2.2.3.

Using all available DCH sequences, the presence of putative recombination sites within the genomes of Taiwan DCH strains was examined using different models (RDP, GENECONV, Bootscan, MaxChi, Chimaera, SiScan, 3Seq, LARD) implemented in RDP5 (version 5.23) [19]. Recombination events with p-values ≤0.05 for at least three different models were considered valid.

## 3. Results

A total of 71 blood samples were analyzed, with 36 (50.70%) coming from female cats, resulting in an equal distribution of sexes within the sample population. The majority (58 out of 71, 81.69%) of the sampled cats were neutered, whereas only about a quarter were exclusively residing indoors (15 out of 71, 21.13%). The age range of the sampled cats spanned from 6 months to 9 years, with a median age of 3 years. Most of these cats (65 out of 71, 91.55%) appeared to be in good health, showing no clinical signs of any disease, while the remaining six were presented with non-specific symptoms, mainly poor appetite.

In the 71 blood samples detected by quantitative real-time polymerase chain reaction, eight (11.27%) were positive for DCH. Four of the eight detected cases were from Kaohsiung City, while two cases each were from Tainan City and Pingtung County (Figure 1). The quantitative estimates of the viral load in these samples varied, ranging from 3×10^5^ to 6×10^8^ copies/mL (Table 1). Furthermore, five of the eight blood samples were tested for selected blood biochemistry markers related to liver function. Out of these five, three (3/5; 60%) of the tested samples had high levels of alanine transaminase (ALT) and aspartate transaminase (AST), but all samples were normal for alkaline phosphatase (ALKP) (Table 2). Likewise, DCH-positive cats were tested for feline immunodeficiency virus (FIV) and feline leukemia virus (FeLV), among them only 1 FIV positive was detected, which is case number 23-05-30-017 (1/4; 25%) and none were detected with FeLV (0/4; 0%) (Table 2).

**Table 2.** Information on the sampled domestic cats positive for domestic cat hepadnavirus.

| Case Number | Age (years) | Sex | Viral Load* (copies/mL) | ALT <sup>1,*</sup> (U/L) | AST <sup>2,*</sup> (U/L) | ALKP <sup>3,*</sup> (U/L) | FIV/FeLV | Strain Code (GenBank Acc. No.) |
| --- | --- | --- | --- | --- | --- | --- | --- | --- |
| 23-05-30-004 | 2 | M | $\sim 3 \times 10^5$ | 400 | 219 | 19 | Negative/<br>Negative | DCH/NPUST-001/TWN/2023 (OR515499) |
| 23-05-30-005 | 1 | M<br>(neutered) | $\sim 7 \times 10^7$ | 41 | 34 | 14 | Negative/<br>Negative | DCH/NPUST-002/TWN/2023 (OR515500) |
| 23-05-30-009 | 1 | F<br>(neutered) | $\sim 7 \times 10^7$ | 170 | 78 | 19 | - | DCH/NPUST-003/TWN/2023 (OR515501) |
| 23-05-30-017 | 2 | M | $\sim 4 \times 10^7$ | 48 | 32 | 75 | Positive/<br>Negative | - |
| 23-05-30-019 | 4 | F<br>(neutered) | $\sim 4 \times 10^5$ | 232 | 62 | 36 | Negative/<br>Negative | DCH/NPUST-004/TWN/2023 (OR515502) |
| 23-06-22-005 | 4 | F<br>(neutered) | $\sim 3 \times 10^8$ | - | - | - | - | DCH/NPUST-005/TWN/2023 (OR515503) |
| 23-06-22-008 | 5 | F<br>(neutered) | $\sim 3 \times 10^6$ | - | - | - | - | - |
| 23-06-22-014 | - | - | $\sim 6 \times 10^8$ | - | - | - | - | DCH/NPUST-006/TWN/2023 (OR515504) |

\* Measured at first sampling. Normal Range: <sup>1</sup> 12-130 U/L; <sup>2</sup> 0-48 U/L; <sup>3</sup> 14-111 U/L. Values outside the normal range are indicated in bold. (-) information not available, or test not performed.

**Figure 1.**
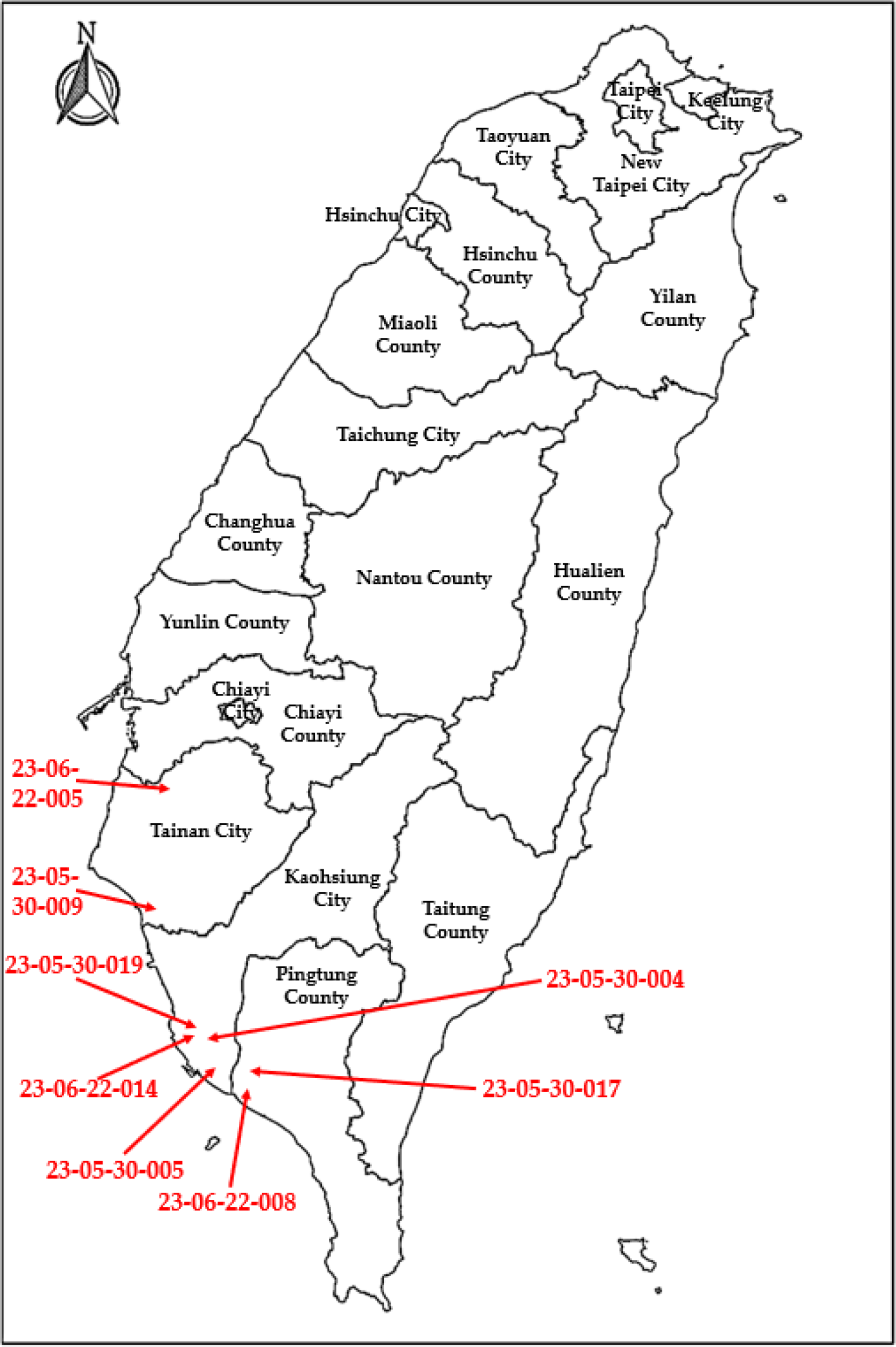
Geographical location of the DCH-positive cats (red arrow). The Taiwan map photo is sourced from the National Land Surveying and Mapping Center (NLSC).

We were able to follow through on four of the five cases 2.5 months after the DCH detection. Viral load and liver health markers were remeasured; the results are shown in Figure 2. All the cats had increased viral load relative to the initial measurement at Day 0. Except for case 23-05-30-004, three of the four cats had increased AST levels; two cases (cases 23-05-30-005 and 23-05-30-019), which were previously within the normal AST range, were found to have abnormal AST levels during the follow-up. Meanwhile, ALT levels for two cases (case 23-05-30-004 and 23-05-30-017) decreased, with the latter going back within the normal range. The other two cases on the other hand were observed to have increased ALT levels, with case 23-05-30-005, which was previously in the normal ALT range, observed to be outside the normal ALT range during follow-up. All cats still tested normal for ALKP 2.5 months after positive DCH detection.

**Figure 2.**
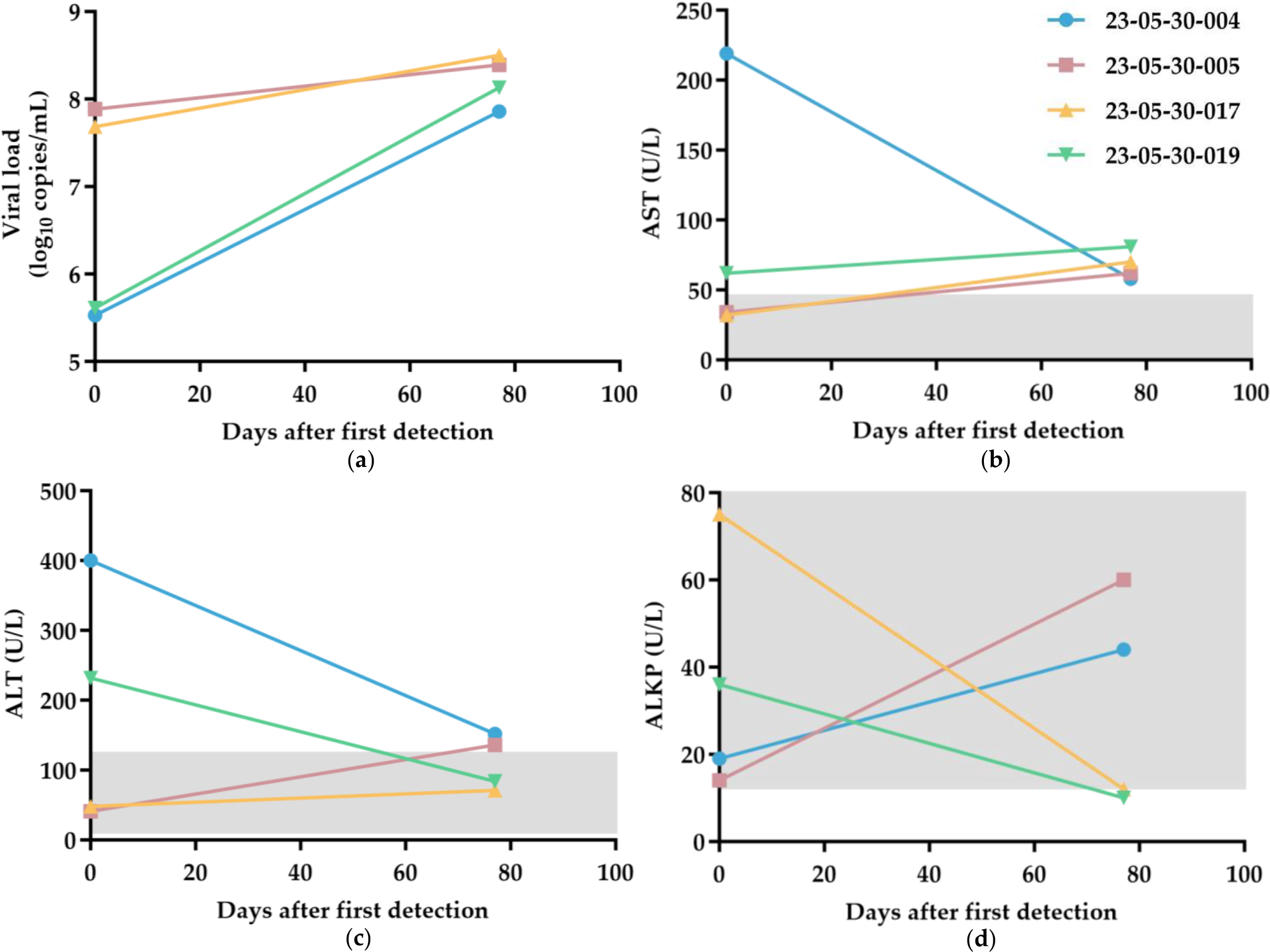
Changes in the (**a**) viral load, (**b**) AST, (**c**) ALT, and (**d**) ALKP levels of DCH-positive cats 2.5 months after first detection. Values in the graph within the normal range for **b**-**d** are indicated by the grey shade. Normal Range: AST 0-48 U/L; ALT 12-130 U/L; ALKP 14-111 U/L.

For further analysis, total DNA was extracted from six selected sera from the positive blood samples. The whole genome of DCH was amplified using three primer pairs. Purified PCR amplicons were quantified by nanodrop, and the three amplicons from a single sample were combined in equal amounts and processed for Oxford Nanopore Technologies sequencing following the rapid barcoding kit protocol (SQK-RBK114.24). A total of more than 150,000 reads (N50 = 933 b) were generated, of which more than 100 Mb were retained after adapter trimming.

Complete closed genomes were successfully amplified and sequenced from six blood samples. The genomes were 3,184 bp in length. Homology analysis using BLASTn search against the NCBI nr database revealed that, at the full genome level, five Taiwan strains (DCH/NPUST-001/TWN/2023 to DCH/NPUST-005/TWN/2023) showed the highest similarity of about 98.7% to DCH strain TR-404 from Türkiye, while another strain (DCH/NPUST-006/TWN/2023) had the highest similarity to PK98-B/THA/2022 from Thailand with a percent ID of 99.1%. Besides that, Taiwan strains shared 97.5% to 100% nucleotide pairwise identities with each other. These observations together suggest the presence of multiple and diverse strains of the DCH in Taiwan. Meanwhile, no evidence of genetic recombination events was detected in the genomes of the Taiwan strains when queried together with all other available DCH genome sequences.

Our phylogenetic analysis using the maximum likelihood method on all available complete sequences of DCH, and genomes of other members of the Orthohepadnavirus, Metahepadnavirus, Herpetohepadnavirus, and Avihepadvirus genera supports previous observations reporting clustering of the available sequences broadly according to their genera (Data/ML tree not shown), as previously reported[5]. On the other hand, among the sequences of DCH, DCH/NPUST-006/TWN/2023 strain grouped with Hong Kong (HK03/2020/16) and Thai (KB18-B/THA/2022 & PK98-B/THA/2022) strains forming a distinct phylogenetic lineage that is sister to the clade where the prototypic Sydney2016 strain belongs (Figure 3). The other Taiwan strains, meanwhile, formed a distinct clade occupied only by the sequences from this study and sister to the clade where most of the genomes recovered from Türkiye belong.

**Figure 3.**
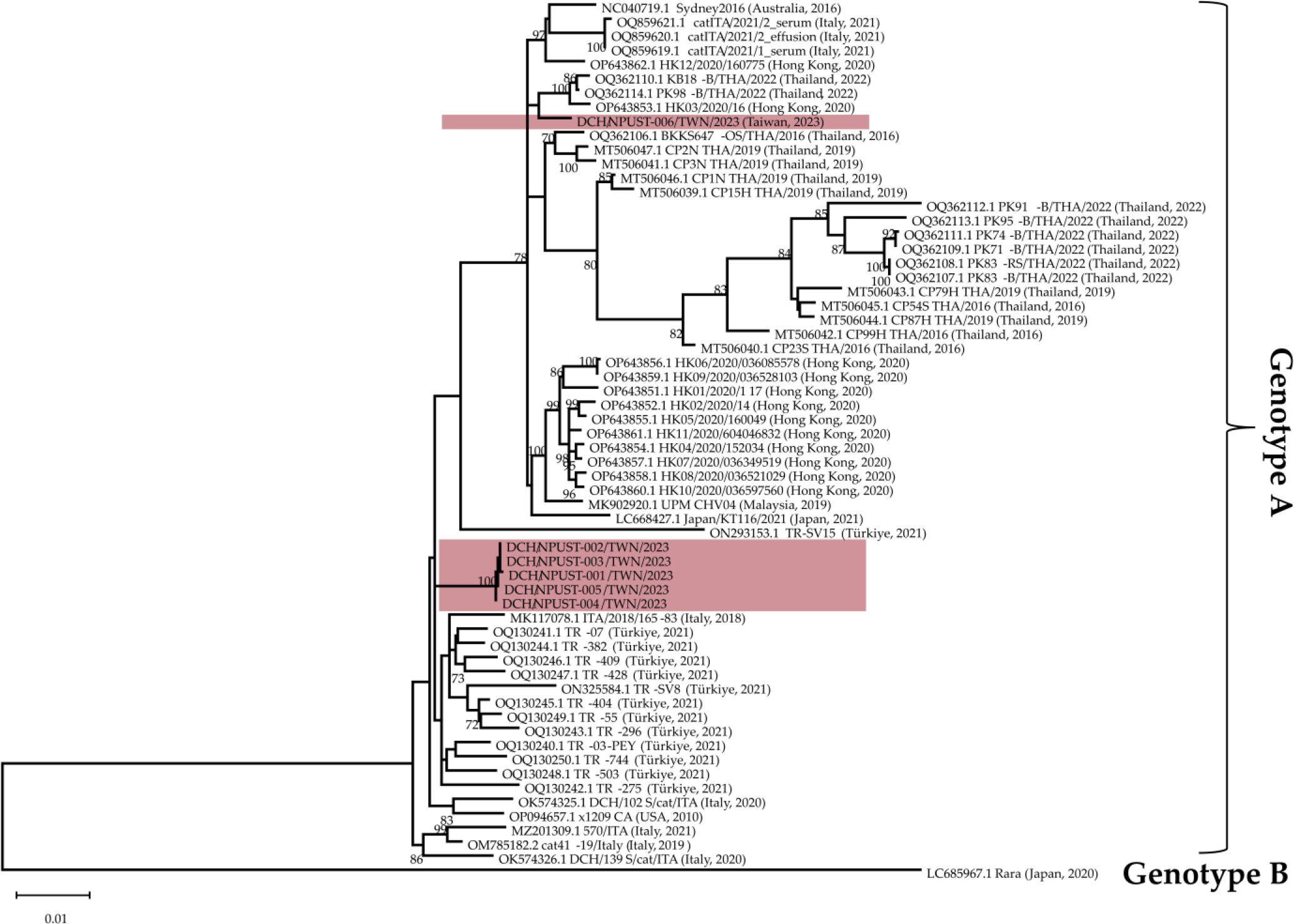
The phylogenetic tree was constructed using complete genomes of DCH collected from Taiwan and available sequences from GenBank. The tree was created by constructing a maximum likelihood tree with TIM+F+I+R2 as the best fit and chosen model according to the Bayesian information criterion using IQTREE v.2.2.2.3. Bootstrap values above 70% are shown. The scale bar indicates nucleotide substitution per site.

Generally, DCH/NPUST-006/TWN/2023 belongs to clade A1, proposed previously by other authors [12], while the rest of the Taiwan strains belong to clade A2 under genotype A. However, with the addition of more sequences particularly those from Türkiye and Taiwan strains, our phylogenetic analysis of the currently available DCH sequences, implemented by constructing a maximum likelihood tree using TIM+F+I+R2 as the best fit and chosen model according to the Bayesian information criterion, did not reveal robust branching support for a clear distinction between the proposed clades A1 and A2 (Figure 3). Rara strain from Japan, however, remains the only member of DCH genotype B.

Pairwise identity scores using all available sequences of the four DCH proteins revealed the highest conservation in core protein ranging from 97.7 to 100 percent ID. On the other hand, X protein showed the greatest variation with percent IDs ranging from 75.2 to 100. The DCH strains’ polymerase and surface protein percent IDs ranged from 85.4 to 100. Phylogenetic relationships among the Taiwan strains, particularly the divergence of DCH/NPUST-006 from the other strains as observed in Figure 3, are strongly supported by the phylogenetic trees based on sequences of core, polymerase, surface, and X proteins (Figure 4).

**Figure 4.**
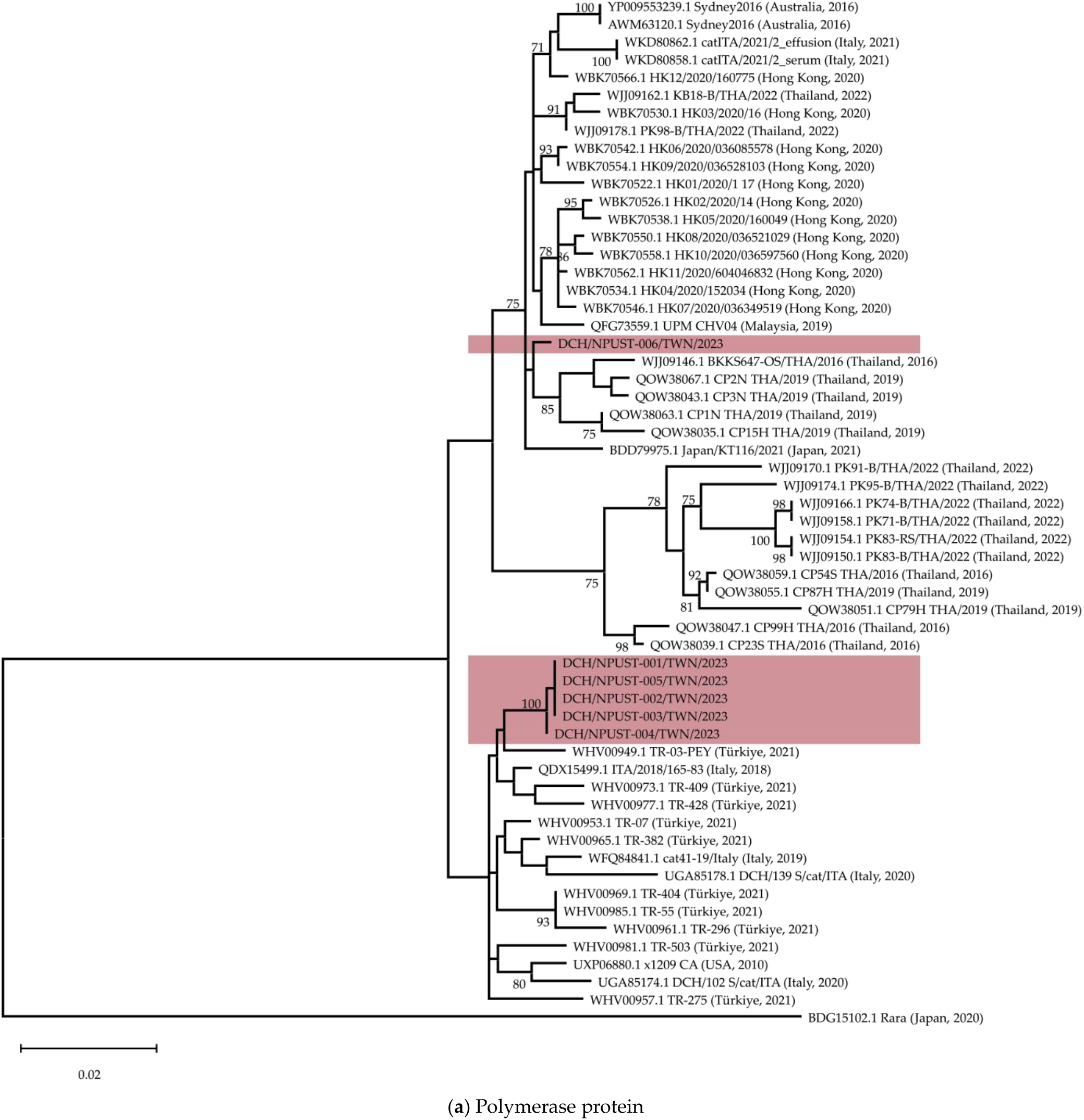
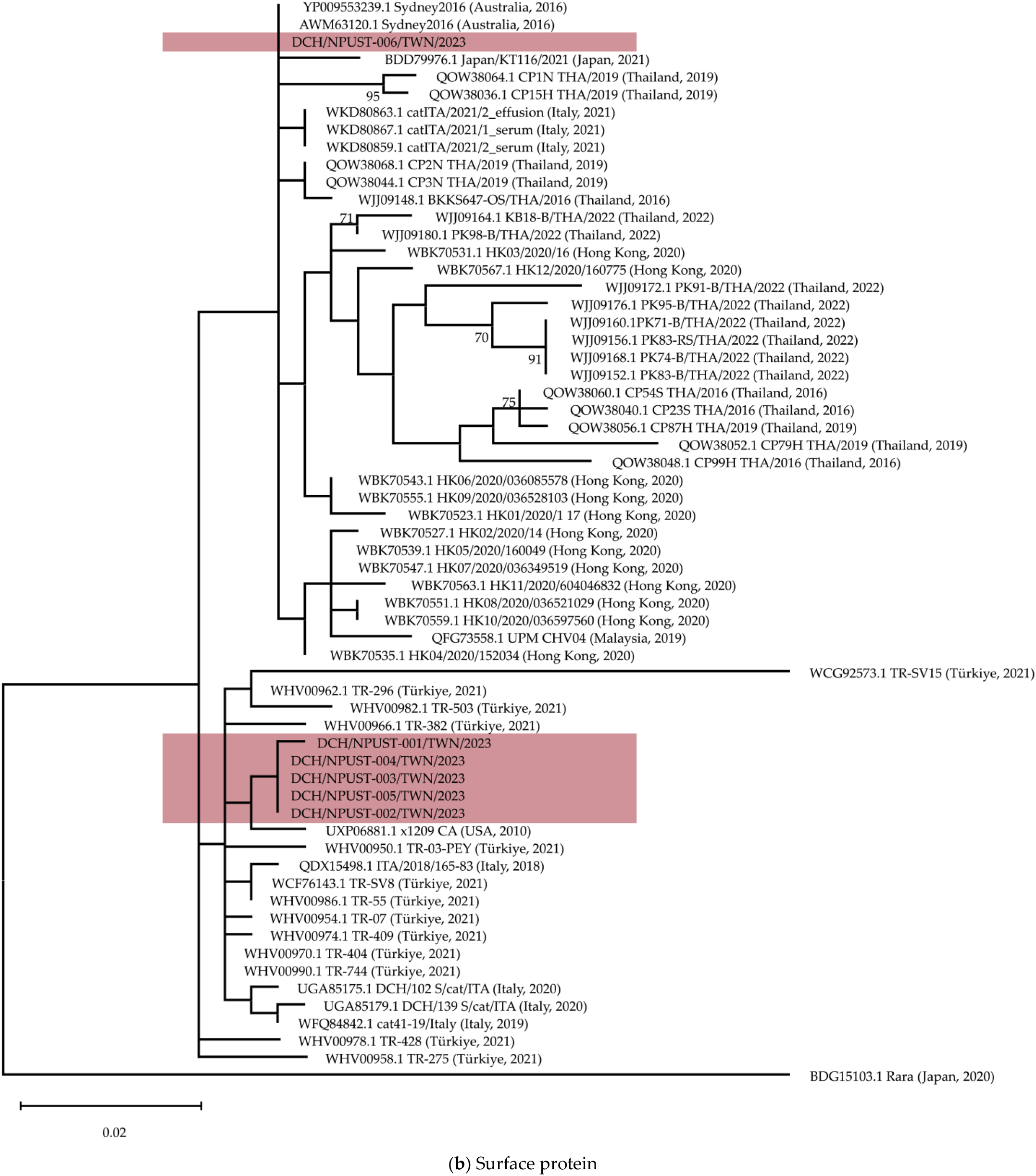
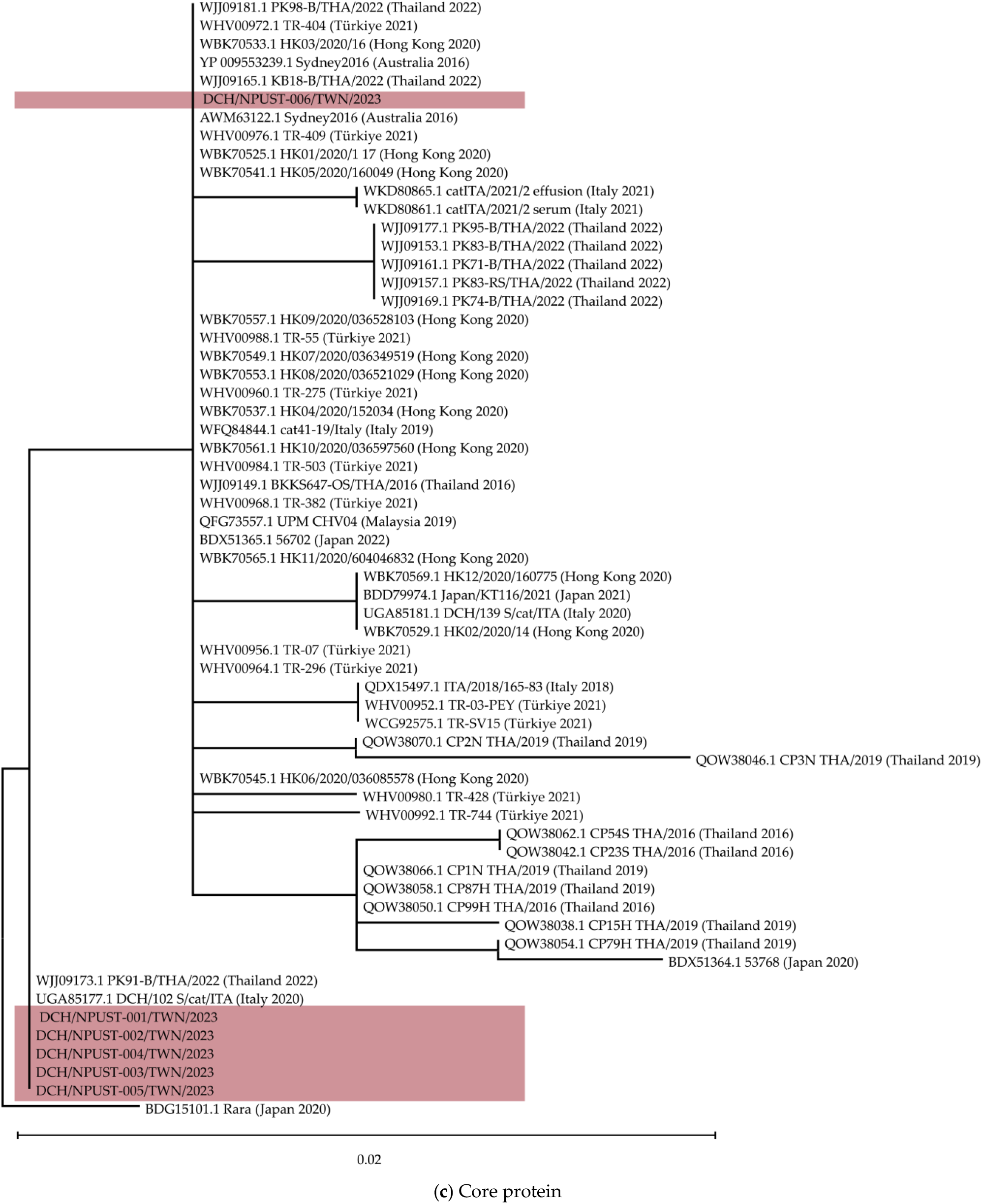
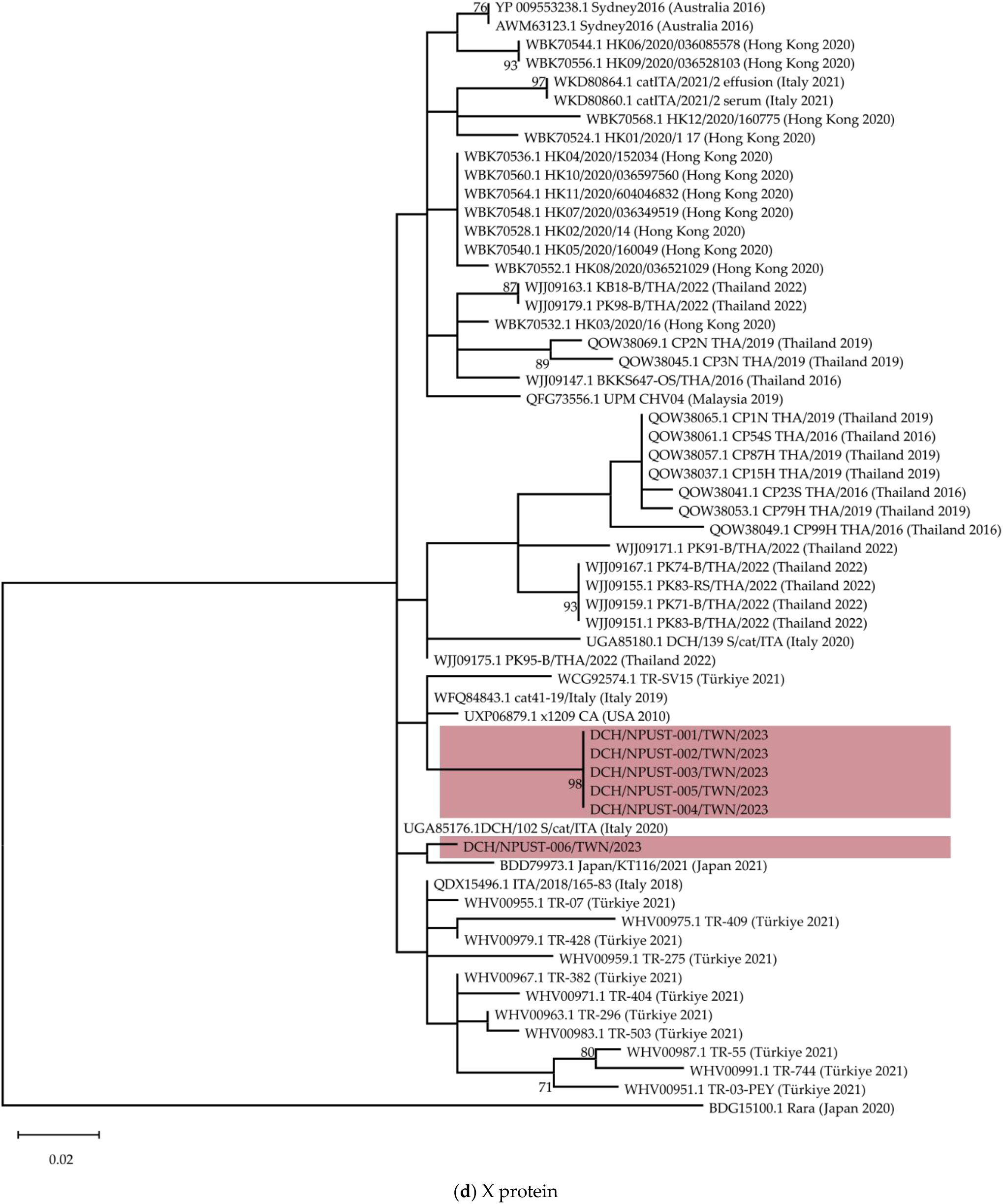
Phylogenetic trees were constructed using the protein sequences for (**a**) polymerase, (**b**) surface, (**c**) core, and (**d**) X proteins of DCH strains from Taiwan and available sequences from GenBank. The tree was created by constructing a maximum likelihood tree with the best fit and chosen model according to the Bayesian information criterion using IQTREE v.2.2.2.3. Bootstrap values above 70% are shown. The scale bar indicates the number of substitutions per site.

Maximum-likelihood trees of both the polymerase and surface proteins (Figures 4a and 4b) generally show the two subdivisions of Genotype A as proposed previously by other authors [12]. However, similar to that of the whole genome sequence tree, there is no robust bootstrap support for this proposed subdivision. Furthermore, such a distinct division is also not apparent in the phylogenetic trees based on the core and X proteins (Figure 4c and 4d). Of note however, strains DCH/NPUST-001/TWN/2023 to DCH/NPUST-005/TWN/2023 were separately grouped with strain each from Italy and Thailand in a clade basal from all other Genotype A strains (Figure 4c).

## 4. Discussion

In recent years, an increasing number of countries have reported the detection of DCH, a novel infectious viral agent of domestic cats. To elucidate the presence and prevalence of DCH in Taiwan, we screened blood samples from 71 cats and demonstrated that eight (11.27%) cats were DCH-positive. Despite a limited sample number, this study revealed the presence of DCH in Taiwan for the first time. Furthermore, our phylogenetic analyses demonstrated that DCH strains in Taiwan represent a distinct lineage compared with previously reported sequences. These findings suggest the importance of continuous surveillance of DCH and further research to elucidate the pathophysiology and transmission route of DCH.

Consistent with other previous reports, our observations in this study suggest no significant association between sex and DCH detection [4,5]. On the other hand, it is interesting to note that while an earlier study observed a higher DCH detection rate among older cats (ages 5-9 years), only one (1/6, 16.7%) of the detected cases in this study was from the same age group, but the majority of the DCH-positive cases were detected among adult cats (>2 years), which is consistent with previous reports [5,17]. Furthermore, all positive cases were from cats that were allowed access outside of the house or are not exclusively residing indoors, thus opening a possibility of transmission from other cats. It has been previously hypothesized that horizontal transmission of DCH may happen through exposure to contaminated blood or fecal material [3,6,7]. In terms of health status, DCH infection was not detected among cats presenting symptoms of ill condition.

At Day 0, all DCH-positive cats in this study were healthy and displayed no disease or infection symptoms. However, by previous reports, the levels of ALT and AST were elevated in about half (3/5, 60%) of the samples tested for markers of liver condition [4,5]. Virus titers of the samples with elevated hepatic markers are >10^4^ genomic copies/mL, indicative of acute infection or active chronic stages of the disease if HBV titer threshold in humans is applied in DCH infections [4]. However, the samples with elevated markers in this study do not correspond to higher virus titers. Potentially complicating this even further, all samples had virus titers >10^4^ genomic copies/mL threshold, but two (2/5, 40%) had normal hepatic markers. Taken together, the current study echoes the recommendation that DCH be included in veterinary diagnostic panels, particularly in cases clinically suspected of hepatic diseases and in screening for blood donors [4,11].

During follow-up 2.5 months after detection, changes in AST and ALT similarly did not correlate with the change in viral load. Although the available information showed that AST tends to increase with viral load, case 23-05-30-004 proved to be an exception that was observed to have the largest decrease in AST levels even approaching the normal range while also having a significantly increased viral load.

A positive correlation of DCH with FIV and FeLV was previously reported in the literature [3,6]. In this study, among DCH-positive cats, one was positive for FIV (1/4, 25%) while none tested positive for FeLV (0/4, 0%). Considering the few DCH-positive sampling sizes, further extensive detection across Taiwan is needed to further investigate the co-infection of FIV and FeLV with DCH. Likewise, collecting blood samples from cats for detection and continuous monitoring of the virus is challenging. Furthermore, the need for wide-range detection of DCH and co-infection with FIV and FeLV is necessary to evaluate its emerging threat to cats in Taiwan.

Lastly, the genetic diversity observed in this study also suggests the need to continuously surveil this virus among the cat populations in the country and to monitor for the introduction of new strains or the potential emergence of recombinants, as observed in another report [3]. Similarly, more studies must be conducted to establish the pathobiology of the virus and the pathophysiology of the infection. This is of utmost importance considering that DCH and HBV share a significant similarity in cell entry receptors, both utilizing the sodium/bile acid cotransporter (NTCP), and that DCH-derived preS1 peptide binds to NTCPs derived from a broad range of animal species, including humans, which both suggest the potential for interspecies and even zoonotic transmission [20].

## Author Contributions

Conceptualization, K.-P.C., A.S. and J.-P.C.; methodology, A.S., A.D.M., B.B.I.S.; formal analysis, B.B.I.S., and A.D.M.; investigation, B.B.I.S., J.-Y.C., Z.-Y.L., H.-H.Z., H.M., M.S., B.H.A.V., and H.-Y.H.; resources, K.P.C, and A.S.; data curation, B.B.I.S., J.-Y.C., Z.-Y.L., H.-H.Z., and B.H.A.V.; writing—original draft preparation, B.B.I.S.,K.P.C; writing—review and editing, B.B.I.S., J.-Y.C., Z.-Y.L., H.-H.Z., A.D.M., M.S., B.H.A.V., H.M., J.-P.C, H.-Y.H, A.S., K.-P.C.; visualization, B.B.I.S. and B.H.A.V.; supervision, K.-P.C., A.S., A.D.M.; funding acquisition, K.-P.C. All authors have read and agreed to the published version of the manuscript.

## Institutional Review Board Statement

The study was conducted following the protocols and guidelines as approved by the Institutional Animal Care and Use Committee of the National Pingtung University of Science and Technology (NPUST-AICUC), with approval number NPUST-112-079.

## Data Availability Statement

All data generated and analyzed in this study are included in this article.

## Acknowledgments

The authors thank Erika P. Butlertanaka, Yuri L. Tanaka, and Tomoko Nishiuchi for their assistance.

## Disclaimer/Publisher’s Note

The statements, opinions and data contained in all publications are solely those of the individual author(s) and contributor(s) and not of MDPI and/or the editor(s). MDPI and/or the editor(s) disclaim responsibility for any injury to people or property resulting from any ideas, methods, instructions, or products referred to in the content.

